# Temporal transcriptomic and microbiome changes in American bison during experimental SARS-CoV-2 challenge

**DOI:** 10.64898/2026.06.08.730980

**Authors:** Bruna Petry, Paola M. Boggiatto, Alexandra Buckley, Eric D. Cassmann, Kaitlyn Sarlo Davila, Steven C. Olsen, Luis Guilherme Virgilio Fernandes, Bienvenido Tibbs-Cortes, Faith M. Rahic-Seggerman, Mitchell V. Palmer, Ellie J. Putz

## Abstract

SARS-CoV-2 continues to pose a threat to humans as well as domestic and wild animals. The variability in severity of clinical signs, the zoonotic potential, and the host-specific response to infection contribute to the persistence of circulation of disease. In wildlife species white-tailed deer have been shown to be more permissive to infection than bovids. However, amongst bovids, American bison have shown a greater susceptibility than cattle. In this study we investigate the transcriptomic response to experimental SARS-CoV-2 infection in bison over time. Substantial numbers of differentially expressed genes were identified between pre- and 2, 5, 7, 14, and 21 days post-infection. KEGG and GO term analysis identified associations with immune response, inflammatory response, and viral infection including COVID-19. IPA analysis of the SARS coronavirus pathway highlighted differences in signaling at days 2 versus 21 post-infection. We additionally examined changes in the nasal microbiome of bison over the course of experimental infection, which suggested an increase in opportunity for secondary infection causing pathogens such as *Mannheimia*. Collectively this study presents a profile of bison transcriptomic response to SARS-CoV-2 infection and continues to expand our understanding of variation in host response.

**Summary:** SARS-CoV-2 remains a threat to humans, domestic animals, and wildlife. Among bovids, American bison show greater susceptibility than cattle. We characterized the bison transcriptomic response to experimental infection across six timepoints, identifying extensive differential gene expression associated with immune, inflammatory, and antiviral pathways. KEGG, GO, and IPA analyses revealed activation of coronavirus-related signaling and shifts between days 2 and 21. We additionally examined changes in the nasal microbiome of bison over the course of experimental infection, which suggested an increase in opportunity for secondary infection causing pathogens such as *Mannheimia*. The results refine understanding of host responses to SARS-CoV-2 in bison.

## Introduction

Severe acute respiratory syndrome coronavirus 2 (SARS-CoV-2) is the virus responsible for the COVID-19 pandemic, which to date has resulted in over 778 million human cases worldwide [1, 2]. In addition to the global impact on humans, the virus has variable infectivity amongst domestic and wildlife species [3]. The list of species with known susceptibility to SARS-CoV-2 has been steadily increasing over the years following the first official report of SARS-CoV-2 in a dog in February 2021. According to Situation Report 22 from the World Organization for Animal Health (WOAH), by 2023 the number of different species infected with the virus was 29, occurring in 36 different countries, with a total of 775 animal outbreaks worldwide [4]. This includes wildlife hosts such as white-tailed deer (*Odocoileus virginianus*) and mule deer (*Odocoileus hemionus*).

Among large animal wildlife species, cervids such as white-tailed deer have shown increased permissiveness to SARS-CoV-2 infection. White-tailed deer are highly susceptible to virus infection both under experimental [5] and field conditions [6], and their SARS-CoV-2 angiotensin-converting enzyme 2 (*ACE2*) receptor sequence shares a high level of homology with that of humans [7]. In experimental models, white-tailed deer are able to transmit the virus to other deer through days 3-5 post-infection (pi), they develop neutralizing antibodies against the virus as early as day 7 pi, and viral RNA was shown to persist in different tissues for at least 21 days after challenge [5, 8]. In contrast, while no evidence of seroconversion has been found in free-ranging elk species, under experimental infection conditions, American elk (*Cervus elaphus canadensis*) can develop a neutralizing antibody response to infection as early as day 7 pi, which was sustained through day 21 pi [9]. Furthermore, blood transcriptome profile of adult and elk calves experimentally challenged with SARS-CoV-2, demonstrated differentially expressed genes (DEG) related to viral response, immune activation, and antibody production. Further, many DEGs were associated with the Coronavirus Pathogenesis Pathways, specifically during peak viral infection on days 2 and 5 pi, for both adult and elk calves [10].

Compared to cervids, bovid species such as cattle have shown less susceptibility to SARS-CoV-2 infection [11]. However, amongst bovids, North American bison (*Bison bison*) appear to be more permissive to infection than cattle [12, 13]. American bison are an important livestock and wildlife species in the United States and are known to be susceptible to diseases such as brucellosis [14, 15], paratuberculosis [16], and some types of herpesvirus such as Ovine Herpesvirus-2 [17, 18]. It is already known that American bison, European bison (*Bison bonasus*), and buffalos (*Bubalus bubalis*) can be infected with coronavirus species, such as bovine coronavirus (BCoV) [12, 19, 20]. Recently, we and others have shown that bison are also susceptible to SARS-CoV-2 infection [12, 13]. Under experimental conditions, bison did not develop clinical signs after challenge; however, they were permissive to infection, as they developed neutralizing antibodies, showed oronasal viral RNA shedding, developed limited interstitial pneumonia, and had viral RNA persist in lymphoid tissues and lung through 21 days pi [13].

The aim of this study was to investigate the whole blood transcriptomic profile of American bison experimentally challenged with SARS-CoV-2. In addition to evaluating the host gene expression, we also examined the nasal microbiota of bison. The respiratory microbiota is known to play a role in pulmonary immunity [21]. Studies have examined the microbiota in COVID-19 patients, documenting perturbations and dysbiosis [22]; however, few studies have examined the microbiota of animals infected with SARS-CoV-2.

## Materials and Methods

### SARS-CoV-2 challenge, sample collection, RNA isolation, and sequencing

For this study, all procedures and animal work were performed following the approval from the NADC Animal and Care Use Committee (IACUC) and in accordance with the Guide for Care and Use of Laboratory Animals regulations. Nine animals, between 8 and 24 months of age, were accommodated to an agricultural biosafety level 3 (ABSL3) facility. After 25 days acclimation, the animals were intranasally challenged with a viral suspension containing 10^6^ tissue culture infectious dose 50 per ml (TCID_50_/ml) of an ancestral strain of SARS-CoV-2 (USA-WA1/2020), as described in Boggiatto et al. [9] and Palmer et al. [13]. Blood samples were collected on day 0 (control, before challenge for 9 animals), day 2 (9 animals), day 5 (6 animals), day 7 (5 animals), day 14 (5 animals) and day 21 pi (5 animals) (Figure 1), and stored following PAXgene^®^ Blood RNA tube (BD Biosciences, San Jose, CA) recommendations. Sterile cotton swabs were used to swab each of the nares of bison on days 0, 2, 7, 14 and 21 pi. To account for microbial contaminants, control swabs exposed to the air of the enclosure were also collected alongside the nasal swabs on days 2, 7,14, and 21 pi. Swabs were immediately placed in 5 mL of sterile 1X PBS and placed on dry ice for transport to -80 °C storage.

**Figure 1.**
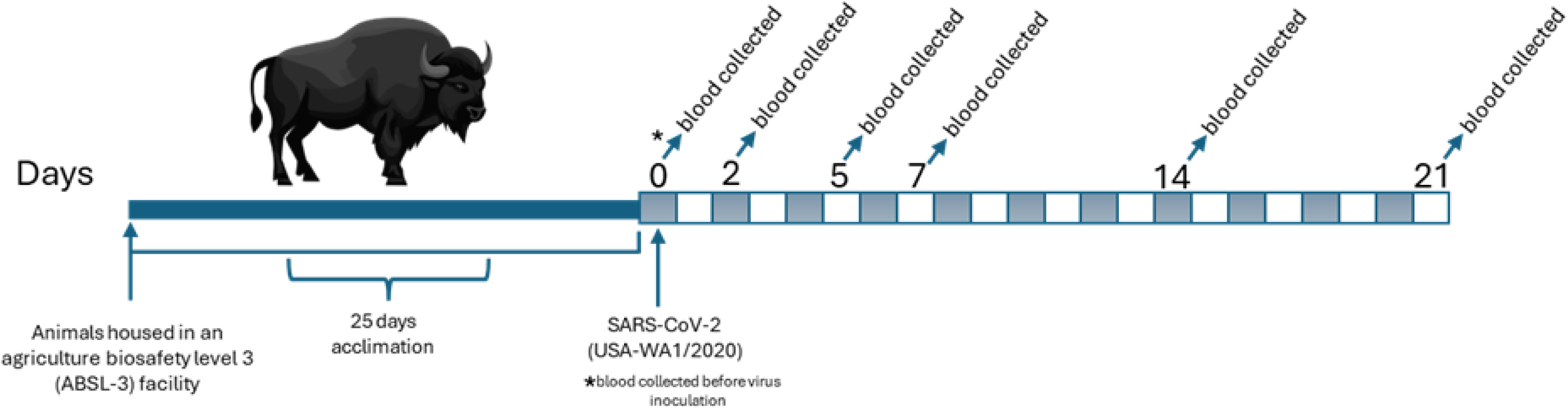
Animals were acclimated in an agriculture biosafety level 3 (ABSL-3) facility for 25 days acclimation, then underwent intranasal SARS-CoV-2 (USA-WA1/2020) challenge. Blood samples were collected on day 0 before virus inoculation (control), 2, 5, 7, 14 and 21 and blood samples were collected.

### RNA isolation, RNA sequencing, bioinformatics: quality control and reads mapping

RNA was isolated using MagMAX™ for Stabilized Blood Tubes RNA Isolation Kit, compatible with PAXgene™ Blood RNA Tubes (catalog number 4451894) (ThermoFisher Scientific, Waltham, MA). The RNA integrity number (RIN) and quality was determined by Bioanalyzer, and samples were submitted to the University of Illinois, Urbana-Champaign, at Roy J. Carver Biotechnology Center for library prep and sequencing. RNA-sequencing was completed using Nova Seq X Plus generating 150 base pair paired end reads.

RNA-sequencing were processed using the Nextflow (version 24.04.2) nf-core/rna-seq pipeline (version 3.14.0) [38, 39]. Read quality was assessed with FastQC (https://www.bioinformatics.babraham.ac.uk/projects/fastqc/, version 0.12.1), followed by adaptor trimming and low-quality read removal (Phred-score<20) using TRIMgalore (https://www.bioinformatics.babraham.ac.uk/projects/trim_galore/, version 0.6.7). After quality filtering, an average of 54 million reads per sample were successfully mapped (mapping quality >0) to the *Bison bison* reference genome (Bison_UMD1.0, GCA_000754665.1) using STAR [29, version 2.7.10a]. Gene-level quantification was performed with featureCounts [30, release 2.0.4]. All the subsequent statistical analyses were performed using R environment (v2024.9.1.394).

### Differentially expressed gene, enrichment analysis and IPA analysis

After the quality control filters, some samples with low quality reads (i.e. less than 10M reads per sample) were removed from downstream analysis. The differentially expressed (DE) analysis was performed comparing each post-inoculation timepoint: day 2 (8 samples), day 5 (6 samples), day 7 (5 samples), day 14 (5 samples) and day 21 (3 samples) to the control group at day 0 (9 samples), using DESeq2 software [31, version 1.44.0]. Prior to the DE analysis, genes were filtered to exclude those with no expression (zero reads), very low expression (less than 1 read per sample on average), and rare expression across the samples (genes with read counts that were not present in at least 20% of the samples). Gene ontology (GO) enrichment analysis and Kyoto encyclopedia of genes and genomes (KEGG) pathway analysis were performed using DAVID Bioinformatics Resources (https://david.ncifcrf.gov/tools.jsp) to help interpret and summarize the relevant biological processes and pathways. To identify canonical metabolic pathways predicted to be activated or inhibited, the DE genes were further analyzed using Ingenuity Pathway Analysis (IPA) (QIAGEN, Redwood City, CA, USA).

### DNA extraction and subsequent longitudinal analysis of the bison nasal microbiota

Tubes containing the frozen nasal swabs were thawed, and biological material was released from the swabs by vortexing at 3200 rpm for 5 minutes using a Vortex-Genie 2 with a 15 ml conical adapter (Scientific Industries, Inc.; Bohemia, NY, USA). After vortexing and removing swabs, biological material was pelleted by centrifugation at 4700 × *g* for 3 minutes. DNA was extracted from the pellet using the DNeasy PowerLyzer PowerSoil Kit (Qiagen; Hilden, Germany) following the manufacturer’s instructions and utilizing the vortex method for the bead beating step. Extracted DNA was sent to the University of Illinois Keck Center for 16S rRNA gene amplicon sequencing. Amplification of the 16S rRNA gene V4 region was performed using primers 515F (5′ GTGYCAGCMGCCGCGGTAA 3′) and 806RB (5′ GGACTACNVGGGTWTCTAAT 3′)(https://earthmicrobiome.org/protocols-and-standards/16s/) on the Fluidigm Biomark HD high throughput PCR system. Pooled libraries were sequenced on a MiSeq V2 flow cell (paired end, 2 × 250 cycles).

Demultiplexed raw reads were merged into contigs by mothur (v1.48.0) [29], and low quality contigs containing any ambiguities, possessing homopolymeric regions > 8 bp, and/or with total length < 252 bp were removed. Merged reads which passed these quality metrics were subsequently aligned against the SILVA SSU NR database (v138); reads aligning outside the region to which 97.5% of all reads aligned were discarded. Finally, chimeric sequences were identified using the SILVA.gold database and subsequently removed. The final, cleaned reads were then clustered into operational taxonomic units (OTUs) using *de novo* clustering at a 99% similarity threshold, and taxonomic classification was performed using the SILVA SSU NR database (v138). Representative sequences from the 10,000 most abundant OTUs were collected from the mothur output and further classified using BLASTn (-word_size = 6, - max_hsps = 1, -max_target_seqs = 10) against the entire 16S rRNA RefSeq database (accessed December 3, 2024). OTUs are referred to by their SILVA.gold classification unless otherwise stated.

Results from mothur were organized and processed using the package phyloseq (v1.48.0) [43] in R (v4.4.1). Next, control samples were used to identify likely contaminants via the prevalence method of the package decontam (v1.24.0) [44]. Using decontam, OTUs were assigned a *P* score representing the likelihood of the OTU originating from the nasal microbiome instead of originating from contamination during sample isolation, processing, or sequencing. *P* score distributions were visualized, and OTUs with a *P* score < 0.725 were classified as contaminants and removed from the analysis along with any remaining OTUs represented in the final dataset by < 10 reads.

Alpha diversity was estimated using the Rust implementation of the R package DivNet, divnet-rs (https://github.com/mooreryan/divnet-rs), in order to calculate the Shannon and Gini-Simpson indices for each time-point [45]. Statistical analysis comparing the alpha diversity indices between each time-point was conducted using the R package breakaway (v4.8.4) [46], accounting for time-point as a fixed effect and using the Benjamini-Hochberg multiple testing correction method. Principal coordinate analysis (PCoA) using Bray-Curtis dissimilarity was performed using phyloseq to visualize beta diversity between the microbiota of each animal at each time-point.

The GLIMMIX procedure in SAS (Version 9.4, SAS Inst., Cary, NC) was employed to determine changes in the abundance of individual phyla, genera, and OTUs following a negative binomial distribution [47]. The statistical model included time-point as a fixed effect, while animal ID was specified as a random effect with repeated measures. Given that four bison were sacrificed after 2 days pi, two separate analyses were completed: (1) comparison of pre-infection versus 2 days pi using samples from all bison (n=9), and (2) pairwise comparisons across all time-points from the surviving bison (n=5). Taxa considered in the analyses included all phyla, the 100 most abundant genera, and the 300 most abundant OTUs. The MULTTEST procedure in SAS was used to control for false discovery rate, and results were deemed statistically significant if both *p* ≤ 0.05 and *q* ≤ 0.05 [48]. For taxa exhibiting significant changes in abundance over time, log_2_fold-changes between time-points were calculated and visualized using R (v4.4.1).

## Results

### Differential Gene Expression across experimental infection

We sequenced whole blood samples from bison at pre-challenge (day 0), and 2-, 5-, 7-, 14-, and 21-days pi. After differentially expressed gene analysis, we identified that bison had a robust response to infection.

When comparing day 2 pi to pre-challenge (day 0), 1,016 DEG were found; 445 downregulated and 571 upregulated (Figure 2A, Supplementary Table 1). From the upregulated gene list, 16 upregulated genes were previously associated with the Coronavirus disease – COVID-19 KEGG pathway, including Angiotensin I converting enzyme (*ACE*), Neuropilin 1 (*NRP1*), and Signal transducer and activator of transcription 1 (*STAT1*). Upregulated genes were additionally associated with other virus-related pathways such as Hepatitis C, Influenza A, and Epstein-Barr virus infection (Supplementary Table 2). Seventeen genes were found to be associated with the innate immune response GO term, including Ficolin 1 (*FCN1*), Nucleotide-binding oligomerization domain-containing protein 2 (*NOD2*), Syntaxin 11 (*STX11*), and Cluster of differentiation 14 (*CD14*). In contrast, among the downregulated DEGs, 46 were annotated within the Herpes simplex virus 1 infection KEGG pathway, suggesting a suppression of select antiviral programs, showing an early host response characterized by both activation of innate immunity and modulation of viral defense pathways (Supplementary Table 2).

**Figure 2.**
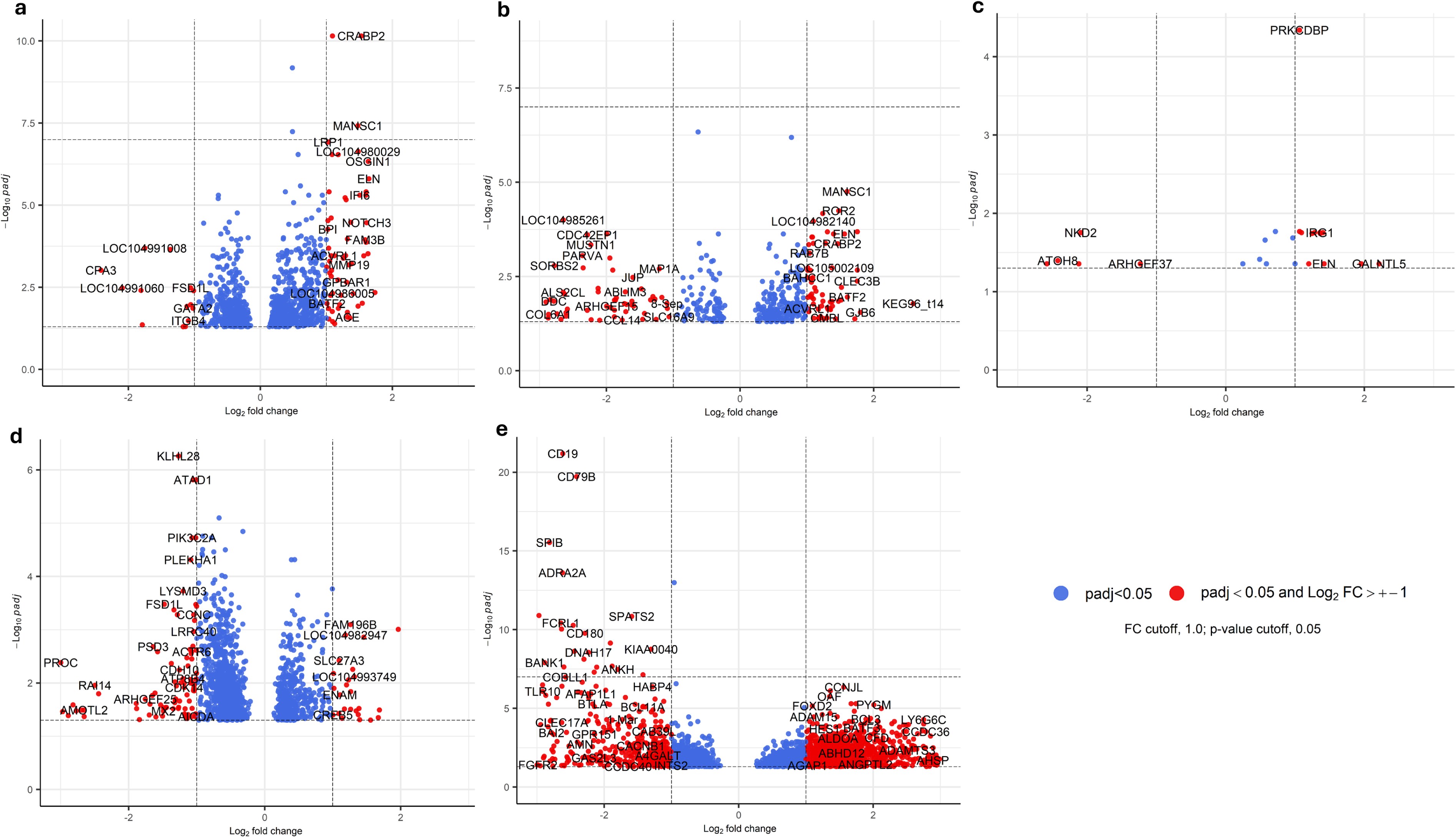
Volcano plots representing differentially expressed (DE) genes with log2-fold changes greater than 1 and significant (p-adjusted < 0.05) for the bison analysis. (a) shows the DE genes on day 2 post-inoculation (pi) compared with those on day 0, (b) for day 5 pi, (c) for day 7 pi, (d) for day 14 pi and, (e) for day 21 pi. Red dots indicate DE genes with an absolute log₂-fold change greater than 1 and the dashed horizontal line represents the adjusted p-value below 0.05. Blue dots are DE genes with p-adjusted < 0.05 and log2-fold change smaller than 1.

Comparing pre-challenge day 0 with day 5 pi, we identified 416 DE genes: 156 downregulated and 260 upregulated (Figure 2B, Supplementary Table 1). Upregulated DE genes were associated with metabolic pathways, lipid and atherosclerosis, cytokine-cytokine receptor interaction and Influenza A, but also GO terms related to immune response and inflammatory responses (Supplementary Table 2). Two examples of the upregulated genes are Interferon induced protein 44 (*IFI44*, log2fc=1.47) and Interferon alpha inducible protein 6 (*IFI6*, log2fc=1.34). *IFI44* plays an important role in the innate immune system during virus infection and can negatively regulate the host viral responses [23, 24]. *IFI6* has already been described during the induction of innate immune responses against influenza A and SARS-CoV-2 [25]. In contrast, the downregulated genes enriched in KEGG pathways included focal adhesion and human cytomegalovirus infection, suggesting possible suppression of cell adhesion mechanisms and viral responses (Supplementary Table 2).

Day 7 pi compared to day 0, yielded only 26 DE genes (Figure 2C, Supplementary Table 1). Of those, 22 genes were upregulated including high expression differences with large log2 fold change values such as *ADRB3* (log2fc = 4.55), *MARCO* (log2fc = 3.98), *TGM1* (log2fc = 3.26), *GALNTL5* (log2fc = 2.21). Of particular interest, macrophage receptor with collagen structure (*MARCO*) is associated with pathogen recognition and inflammatory signaling during infection [26]. Upregulated genes were associated with GO terms including membrane, transmembrane, and WNT-protein binding, highlighting a dynamic shift in cellular communication and immune response pathways (Supplementary Table 2). There were only six DE downregulated genes identified; *FBLN1* (log2fc = -3.10), *ATOH* (log2fc = -2.42), *MUSTN1* (log2fc = -2.12), *NKD2* (log2fc = -2.09) and *ARHGEF* (log2fc = -1.23). Some of these genes are associated with functions including extracellular matrix composition and WNT-signaling, and the reduction in their expression can be reflecting a potential response to infection related cellular stress or immune modulation [27, 28].

Day 14 pi versus day 0 analysis revealed 1,478 DEG; 467 upregulated and 1,011 downregulated (Figure 2D, Supplementary Table 1). Upregulated genes were associated with Endocytosis, Salmonella infection, human T-cell leukemia virus 1 infection, Neurotrophin signaling pathway, and tuberculosis KEGG pathways; GO terms were associated with nucleus, cytoplasm, ATP binding and protein binding. Downregulated DE genes had associations with GO terms such as nucleus, cytoplasm, metal ion binding, protein binding, ATP binding, RNA binding, and cell differentiation, and KEGG pathways predicted associated terms including Herpex simplex 1 virus infection, MAPK signaling pathway, and human papillomavirus (HPV) infection (Supplementary Table 2).

Lastly, day 0 versus day 21 pi, there were 3,078 DE genes: 1,906 upregulated and 1,172 downregulated (Figure 2E, Supplementary Table 1). From the upregulated list, 74 genes were associated with the Coronavirus disease – COVID-19 KEGG pathway including: Innate immune signal transduction adaptor (*MYD88*), NF-κB inhibitor alpha and beta (*NFKBIA* and *NFKBIB*), TNF receptor superfamily member 1A (*TNFRSF1A*) and the mitogen-activated protein kinase 3 (*MAPK3*). Also identified KEGG pathways were 77 genes related to pathways of neurodegeneration, 66 genes in the Alzheimer disease pathway, and 51 genes in the Parkinson disease (Supplementary Table 2). Additionally, 29 upregulated genes were associated with the GO term for inflammatory response. Downregulated genes were associated with KEGG pathways such as Herpes simplex 1 virus infection (51 genes), ribosome biogenesis in eukaryotes (22 genes), human T-cell leukemia virus 1 infection (20 genes) and cell cycle (19 genes) (Supplementary Table 2).

To take advantage of our longitudinal analysis we assessed whether DE genes were conserved across timepoint comparisons. There were three upregulated DE genes across all timepoints: Solute Carrier Family 35 Member E4 (*SLC35E4*), DC-STAMP Domain Containing 2 (*DCST2*) and Zinc finger and BTB domain-containing 7B (*ZBTB7B*). Five downregulated genes overlapped across days 2, 5, 14 and 21 pi comparisons: Solute Carrier Family 16 Member 9 (*SLC16A9*), Zinc Finger AN1-Type Containing 4 (*ZFAND4*), Zinc Finger Protein 547 (*ZNF547*), G-Patch Domain Containing 2 Like (*GPATCH2L*), Nuclear Casein Kinase and Cyclin Dependent Kinase Substrate 1 (*NUCKS1*) (Figure 3).

**Figure 3.**
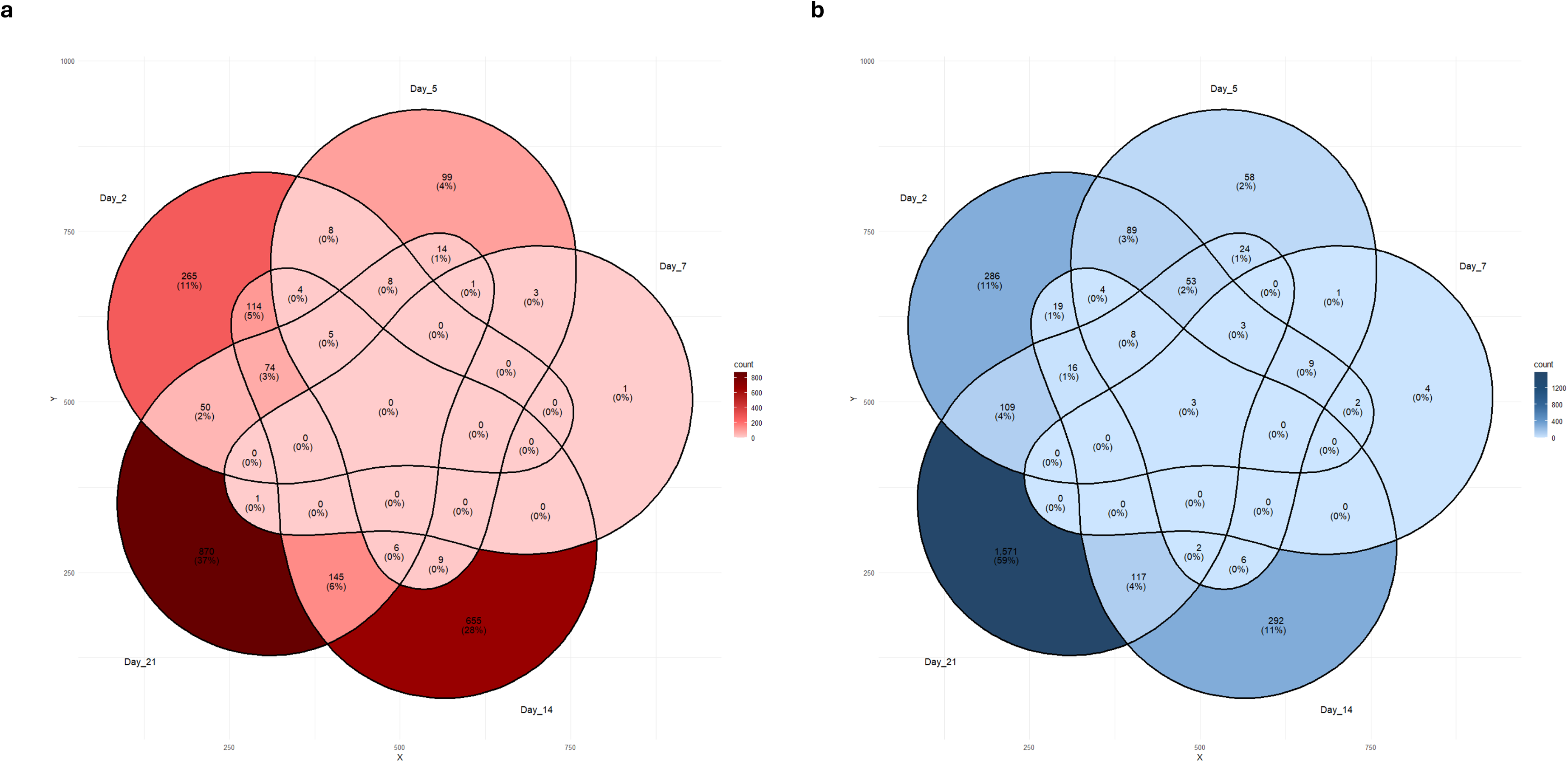
Venn Diagram shows the number of conserved up and downregulated genes in all time points. Figure (a) shows downregulated genes and (b) upregulated genes.

### Ingenuity Pathway Analysis (IPA)

We utilized the IPA software to characterize nuances within the coronavirus disease pathway identified in our differential expression analysis. IPA analysis corroborated KEGG pathway findings and identified associations with DE genes and coronavirus diseases on both days 2 and 21 pi (see Figure 4). Early in infection (day 2 pi) we see C-C motif chemokine receptor 2 (*CCR2*) and Interleukin-17 receptor A (*IL17RA*) have increased expression compared to pre-challenge. Interferon-stimulated gene factor 3 (*ISGF3*) was predicted to be activated compared to day 0 at both days 2 and 21 pi and it is suggested to contribute to the inhibition of SARS-CoV-2 replication and activation of the adaptive immune response. Terms related to innate immunity were also activated at both timepoints through IFN type 1 activation, where a pro inflammatory cytokine response is predicted through NF-kB signaling. Angiotensin II Receptor Type I (*AGTR1*) was activated on day 21 pi and indirectly related to vasoconstriction, acute myocarditis, acute respiratory distress syndrome (ARDS) and inflammation. Also on day 21 pi, SMAD family members 3 and 4 (*SMAD3* and *SMAD4*) were predicted to be inhibited. Unique to the 21-day pi comparison, Interferon-induced transmembrane protein 1 (*IFITM*) was upregulated and indirectly related to endocytosis of virus inhibition, as well as heterogeneous nuclear ribonucleoprotein A1 (*HNRNPA1*) which is also indirectly related to inhibition in the replication of SARS-CoV-2.

**Figure 4.**
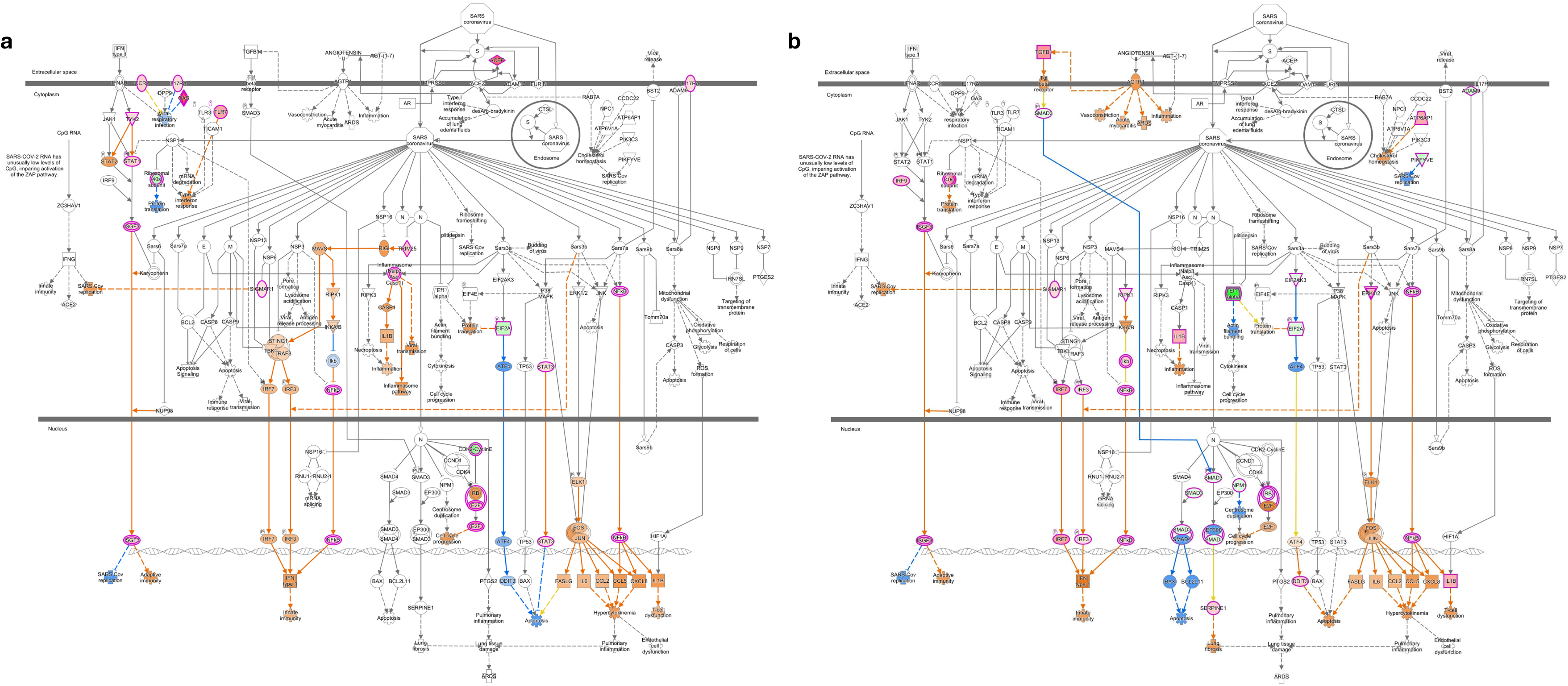
Ingenuity pathway analysis (IPA) showing the differentially expressed (DE) genes involved in the coronavirus pathogenesis pathway in bison. The Core analysis shows predicted activated and inhibited pathway associations for the DE gene lists for day 2 (a) and day 21 (b) post-SARS-CoV-2 infection. Downregulated genes are shown in green, and upregulated genes are shown in pink. The Molecule Activity Predictor tool was used to predict downstream activity based on significant differential gene expression. Predicted activation is shown in orange, and predicted inhibition is shown in blue. The darker the fill is, the more confident the prediction. The solid lines represent direct relationships, whereas the dashed lines represent indirect relationships.

### Longitudinal analysis of the bison nasal microbiome

DNA was isolated from nasal swabs of bison on days 0, 2, 7, 14 and 21 pi. Examination of raw data revealed 9,542,608 merged reads/contigs. Of these, 6,980,907 reads remained following quality control and chimera removal in mothur [29]. *De novo* operational taxonomic unit (OUT) clustering generated 433,194 OTUs of which 296 were classified as likely contaminants; removal of the likely contaminants and low abundance OTUs yielded a final dataset of 29,099 OTUs. Alpha diversity analysis revealed more significant differences in the Shannon index between time-points compared to the Gini-Simpson index. The Shannon index at 2 days pi and 21 days pi was significantly higher than at the pre-infection time point but was significantly lower at 14 days pi. Additionally, alpha diversity was more variable at 14 days pi according to both Shannon and Gini-Simpson indices (Figure 5A). No obvious clustering was apparent from visualization of beta diversity by principal coordinate analysis (PCoA) (Figure 5B).

**Figure 5.**
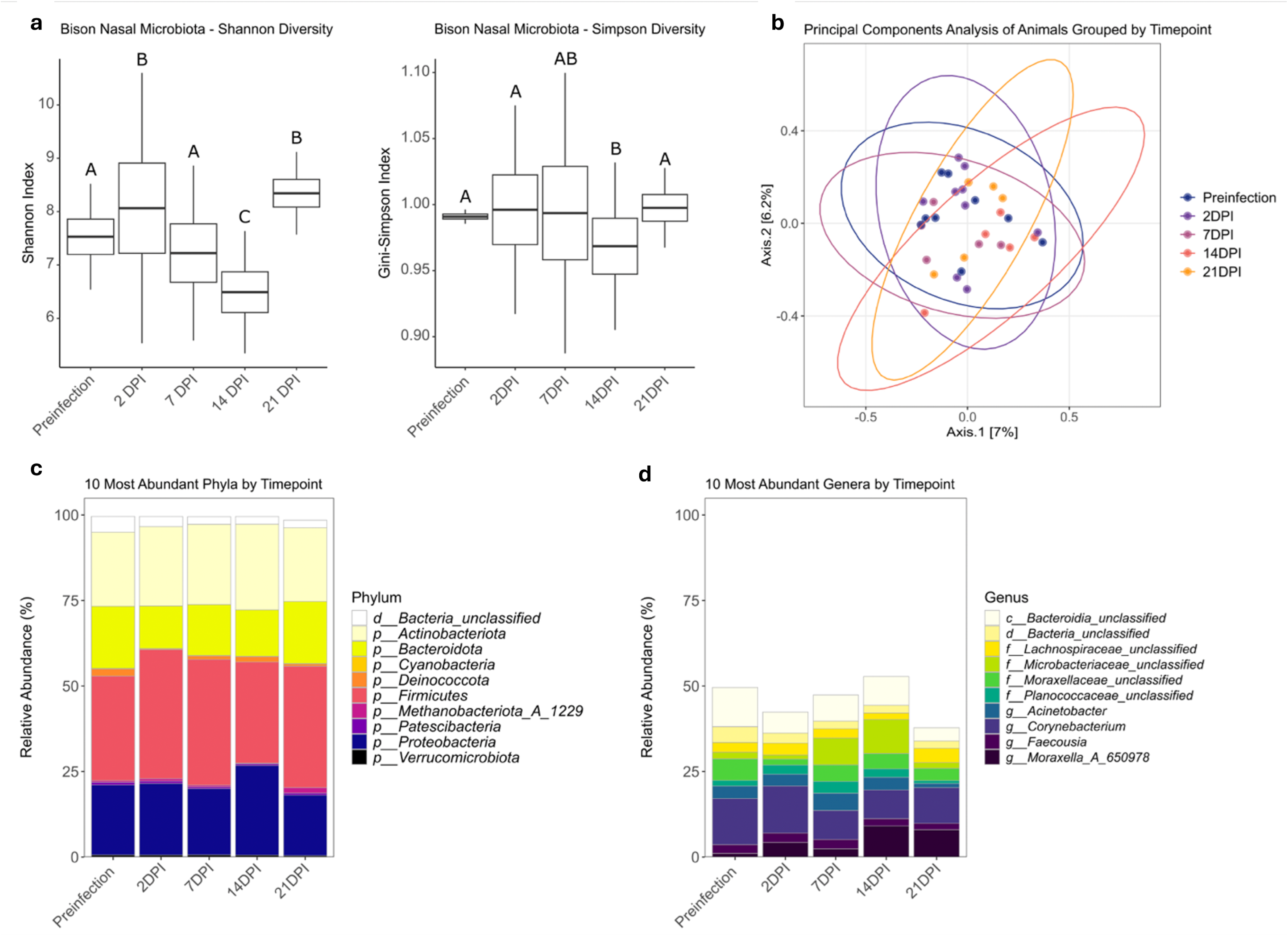
Alpha diversity, beta diversity, and the 10 most abundant phyla and genera of the bison nasal microbiota at all time-points. (A) Shannon (left panel) and Gini-Simpson (right panel) diversity indices of the bison nasal microbiota were calculated using divnet-rs. Time-points with the same letters are not significantly different (*p* > 0.05). (B) Phyloseq was used to conduct principal coordinate analysis (PCoA) based on Bray-Curtis dissimilarity to visualize differences in the composition of the microbiota of each animal at each time-point. Ellipses represent 95% confidence levels assuming a multivariate normal distribution. (C) Relative abundances of the 10 most abundant phyla and (D) genera in the dataset.

The most abundant phyla at all time-points were *Bacillota* (*Firmicutes*), *Pseudomonadota* (*Proteobacteria*), *Actinomycetota* (*Actinobacteria*), and *Bacteroidota* (*Bacteroidetes*) (Figure 5C). Common genera across all time-points included *Corynebacterium*, *Moraxella*, and an unclassified genus in the family *Moraxellaceae* (Figure 5D). Among the most abundant OTUs, OTUs 1 and 2 clustered as part of the same genus and were most similar to *Limnovirga soli* (BLAST percent identity of ∼92%). BLAST analysis also revealed that OTU 3 was most similar to *Candidatus* Limnoluna rubra (∼96% percent identity). OTU 4, which was part of the aforementioned unclassified *Moraxellaceae* genus, was most similar to members of the genus *Psychrobacter* (∼96% percent identity) (Supplementary table 3).

Between the pre-infection nasal samples and 2 days pi, the relative abundance of 8 phyla (Figure 6A, Supplementary File 4) and 22 OTUs differed significantly (Figure 6B, Supplementary File 4). OTUs which were more abundant at 2 days pi included OTUs 12, 122, and 124 which were all classified as *Mannheimia*. When analyzing only the 5 animals which were sampled at all time-points, 14 phyla, 3 genera, and 80 OTUs were found to be differentially abundant across all time points (Supplementary File 4). Multiple of these OTUs were classified as belonging to taxa known to be associated with bovine respiratory disease: *Mannheimia* (OTUs 12, 123, 124, and 418), *Pasturellaceae* (OTUs 52 and 343), and *Mycoplasmopsis* (*Mycoplasma*) (OTUs 41, 273, and 424). With the exceptions of OTUs 41 and 343, these putative respiratory pathogens increased in abundance relative to the pre-infection time-point at 2 and 7 days pi, and some of these taxa maintained a significantly increased abundance out to 14 days pi (Figure 6C).

**Figure 6.**
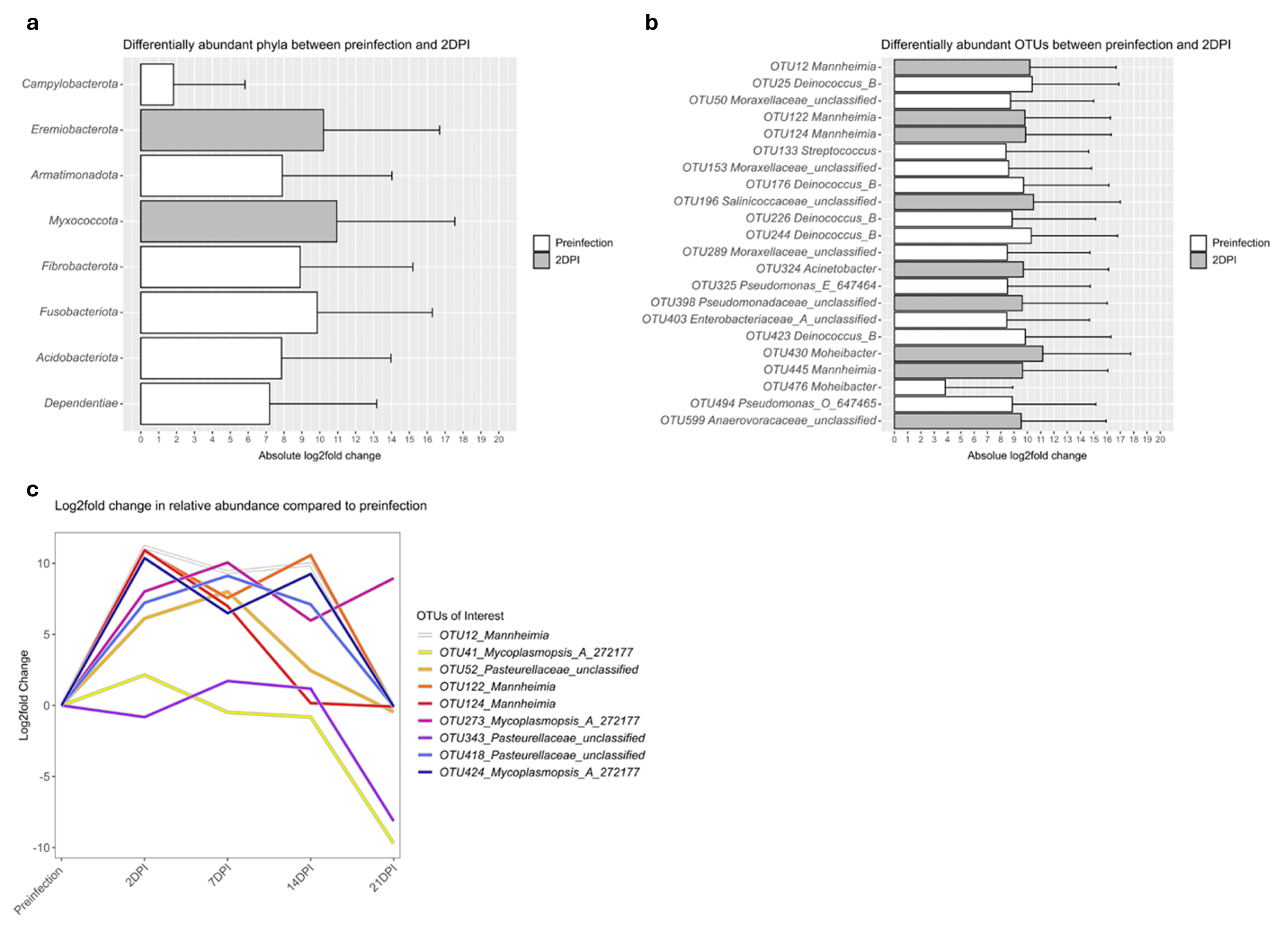
Differentially abundant taxa across time-points. (A) Phyla and (B) genera which are differentially abundant between pre-infection and 2 days pi as expressed by absolute log2fold changes in relative abundance. Data is based on SAS analysis of samples from all 9 animals at pre-infection and 2 days pi (C) Log2fold changes in the relative abundance of selected OTUs at all time-points relative to the relative abundance at the pre-infection time-point. Data is based on SAS analysis of samples from only the 5 animals which were sampled at each time-point.

## Discussion

American bison appear more permissive to infection than their cattle bovid counterparts as illustrated collectively by oral and nasal detection of viral RNA, production of neutralizing antibodies by day 14 pi, the presence of microscopic lesions during the acute disease phase, and consistent viral RNA detection in the medial retropharyngeal lymph nodes and lung throughout infection [13]. To further understand the immune mechanisms driving the bison response to SARS-CoV-2, we investigated the bison blood transcriptomic response to experimental SARS-CoV-2 infection. DEG changes of interest were associated with all timepoints, with an emphasis on the COVID pathway associations identified on days 2 and 21 pi.

Our group is interested in host components that might contribute to the difference in susceptibility amongst different host species. Previously, we published the transcriptomic responses of adult and elk calves to experimental SARS-CoV-2 challenge. Upon examination of collective KEGG and GO term analysis, there are distinctions between species. Unlike elk, bison do not have as many associations with inflammation markers, neurodegenerative disease, or additional viral pathogens. However, comparing the IPA Coronavirus pathogenesis pathway, both bison and elk have similar recognition profiles early in infection. At 2 days pi, bison, similar to elk [10], have predicted type 1 IFN and NF-kB activation, contributing to innate and adaptive immune responses including upregulation of the interleukin-1 beta (*IL1b*), C-X-C motif chemokine ligand 8 (*CXCL8*), C-C motif ligand 5 (*CCL5*), and interleukin-6 (*IL-6*). Notably, in elk we see decreased expression and inhibition of these activating networks and cytokines by day 14 pi, however, in bison there is still strong *NF-kB* and *type 1 IFN* pro-inflammatory signaling even through day 21 pi. Neither elk nor bison displayed clinical signs of disease despite evidence of sustained infection as shown by the persistence of viral RNA in lymphoid tissues through day 21 pi. However, unlike elk, bison also showed persistent viral RNA in lung tissues. Furthermore, the viral RNA in the lungs was associated with pulmonary pathology, specifically focal areas of interstitial pneumonia. Collectively, the bi-modal transcriptomic response observed in bison, with DEG responses first peaking at 2 days pi, a lull at day 7 pi, and subsequent second peak at day 21 pi, is consistent the expected innate and subsequent adaptive immune responses. Furthermore, the extended pro-inflammatory signaling observed in the periphery as indicated by day 21 pi data, along with the pulmonary pathology and viral RNA presence, suggest that bison may be less effective at clearing the virus as compared to elk.

While many factors affect disease pathogenesis, one of these factors may be related to the commensal nasal flora of bison, and disruptions that may occur following an infection. Here, we assessed changes associated with the bison nasal microbiome following infection with SARS-CoV-2. Longitudinal analysis of the bison nasal microbiota during the experiment revealed perturbations in alpha diversity and shifts in the abundance of specific taxa following intranasal SARS-CoV-2 infection. Notably, multiple OTUs belonging to the taxa *Mannheimia*, *Mycoplasmosis* (*Mycoplasma*), and *Pasteurellaceae* were found to increase in relative abundance following SARS-CoV-2 infection relative to the pre-infection time-point. Members of these taxa include opportunistic pathogens of bison and cattle, including some of the causative agents of the bovine respiratory disease complex [30–33]. Bison and cattle are susceptible to *Mannheimia* infection, but *in vitro* work has shown greater reactive oxygen species and bacterial uptake by cattle compared to bison [34], suggesting that cattle have a more robust response to infection with this bacterium. This could suggest that bison, and/or other bovids, may be susceptible to comorbidities or to co-infections in the presence of SARS-CoV-2. An increase in relative abundance of opportunistic respiratory pathogens in the nasal cavity following SARS-CoV-2 infection may have negative implications for bison health even if the animals exhibit a low level of susceptibility to SARS-CoV-2 infection itself. Indeed, a pattern of increased opportunistic pathogens in the human nasopharyngeal microbiota following SARS-CoV-2 infection has also been observed [35–37].

Bison are one of the few bovids that have shown to be permissive to SARS-CoV-2 infection, thus far. Characterizing the variable immune response to SARS-CoV-2 in different host species offers valuable insight into differences in susceptibility, as well as transcriptomic profiles that may contribute to establishing biomarkers or improving surveillance and diagnostics.

## Data availability

Microbiome and RNA sequencing data generated and analyzed during this study are available under Bioproject PRJNA1226260. BioProject and associated SRA metadata are available at “https://www.ncbi.nlm.nih.gov/bioproject/PRJNA1226260”. All other data generated or analyzed during this study are included in this published article (and its Supplementary Information files).

## Acknowledgments

We gratefully acknowledge the National Animal Disease Center (NADC) for their support in animal care throughout this study. In particular, we thank Dr. Rebecca Cox, Derek Vermeer, Jonathan Gardner, Tiffany Williams, Emma Prat, Emma Hay, and Jared Peterson for their dedicated assistance with animal husbandry and care. We also extend our appreciation to Sue Osorio, Shelly Zimmerman, and Sarah Anderson for their excellent technical support. Additionally, we thank the University of Illinois for performing sample sequencing.

## Funding

This research was supported by intramural funding from the U.S. Department of Agriculture and by American Rescue Plan (ARP) funds through an interagency agreement between the Agricultural Research Service (ARS) and the Animal and Plant Health Inspection Service (APHIS) Wildlife Services (WS), and USDA ARS CRIS #3625-32000-232-00D. Additional support was provided through an appointment to the ARS Research Participation Program, administered by the Oak Ridge Institute for Science and Education (ORISE) via an interagency agreement between the U.S. Department of Energy (DOE) and USDA. ORISE is managed by ORAU under DOE contract number DE-SC0014664. The views and opinions expressed in this publication are those of the authors and do not necessarily reflect the official policies or positions of the USDA, DOE, or ORAU/ORISE.

## Competing interests

The authors declare no competing interests.

## Author contributions

B.P. analyzed the data and wrote the manuscript.

P.M.B. participated in study design, performed the animal experiments, collected samples, analyzed data, and reviewed the manuscript.

A.C.B. participated in study design, prepared the virus for infections, assisted with animal work, sample collection, and sample processing.

E.D.C. participated in study design, collected samples.

K.S.D. analyzed data and discussion.

S.C.O. performed animal experiments and collected samples.

B.T.C. sample collection and analyzed microbiome data.

F.M.R.S. sample collection.

M.V.P. obtained funding, managed projects, participated in study design, performed animal experiments, and collected samples.

E.J.P. participated in the study design, processed samples, analyzed data, helped write and reviewed the manuscript.

## Supplementary Material

Supplementary table 1 – List of differentially expressed (DE) genes across all time points following SARS-CoV-2 challenge. DE genes were identified by comparing each challenged group to the uninfected control group (day 0).

Supplementary table 2 – Enrichment analysis of differentially expressed (DE) genes following SARS-CoV-2 challenge. The table includes significantly enriched KEGG pathways and Gene Ontology (GO) terms associated with upregulated and downregulated genes.

Supplementary table 3 – Taxa information from the most abundant OTUs (top 10k) based on BLAST analysis, showing the similarities between member of the same genus.

Supplementary table 4 – Relative abundance of different phyla and OTUs on day 2 post-infection.

---

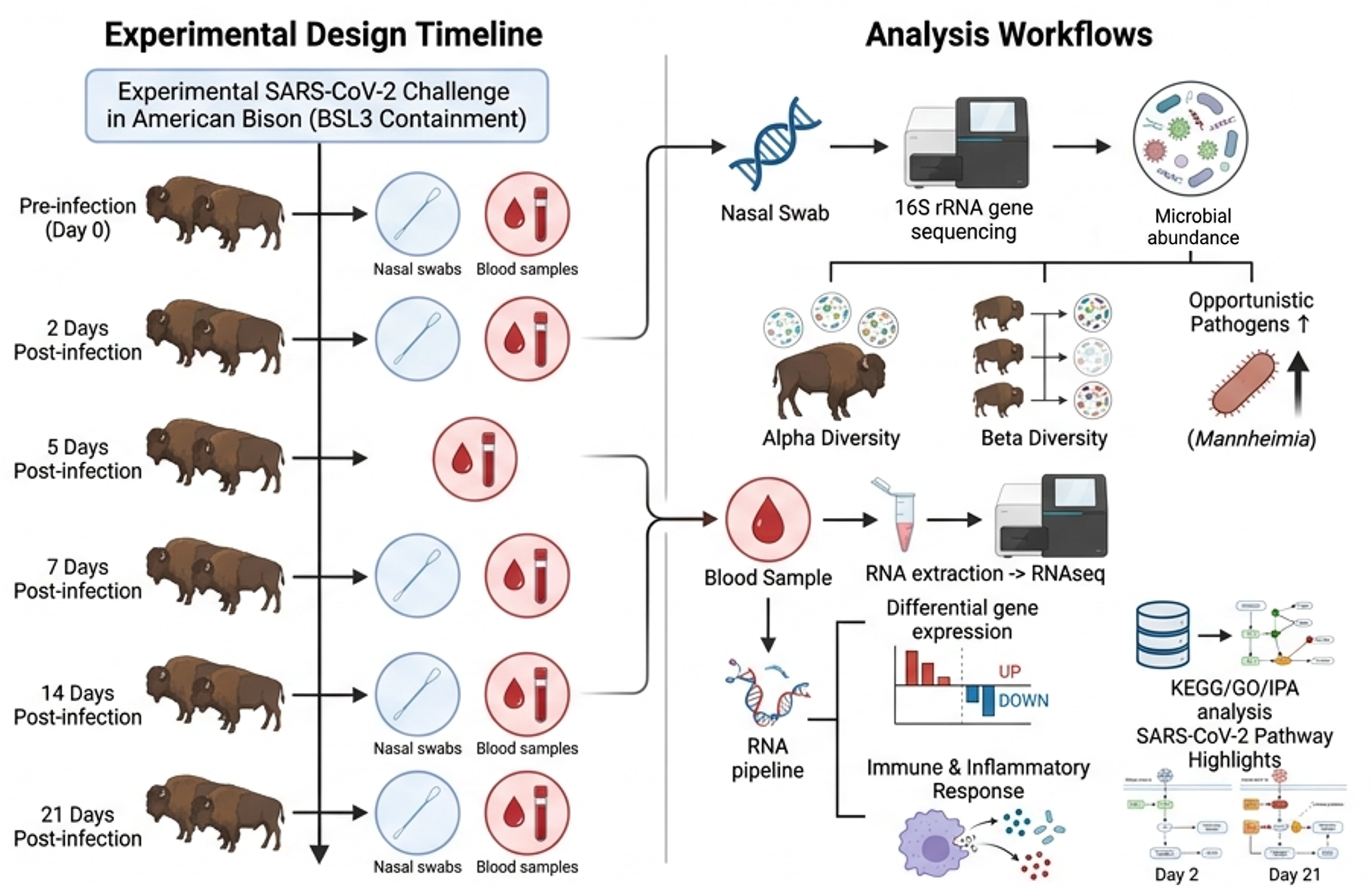

